# A human huntingtin SNP alters post-translational modification and pathogenic proteolysis of the protein causing Huntington disease

**DOI:** 10.1101/129536

**Authors:** DDO Martin, C Kay, JA Collins, YT Nguyen, RA Slama, MR Hayden

## Abstract

Post-translational modifications (PTMs) are key modulators of protein function. Huntington disease (HD) is a dominantly inherited neurodegenerative disorder caused by an expanded CAG trinucleotide repeat in the huntingtin (*HTT*) gene. A spectrum of PTMs have been shown to modify the normal functions of HTT, including proteolysis, phosphorylation and lipidation, but the full contribution of these PTMs to the molecular pathogenesis of HD remains unclear. In this study, we examine all commonly occurring missense mutations in *HTT* to identify potential human modifiers of HTT PTMs relevant to HD biology. We reveal a SNP that modifies post-translational myristoylation of HTT, resulting in downstream alterations to toxic HTT proteolysis in human cells. This is the first SNP shown to functionally modify a PTM in HD and the first validated genetic modifier of post-translational myristoylation. This SNP is a high-priority candidate modifier of HD phenotypes and may illuminate HD biology in human studies.

## Introduction

Huntington disease (HD) is a debilitating neurodegenerative disease with no treatment to delay progression of the disease. HD is caused by an expanded CAG repeat in the huntingtin gene (*HTT*) that translates into a polyglutamine tract at the N-terminus of the huntingtin protein (HTT)^1^. Wild-type HTT is critical for cell viability^2^ and has been shown to play a role in a variety of pathways including orientation of the mitotic spindle^3^, trafficking of autophagosomes in neurons^4^, and in regulating autophagy^5^. Expanded polyglutamine in HTT leads to loss of function and a toxic gain of function in neurons of the striatum and cortex, causing behavioral changes, movement deficits and ultimately death. Age of disease onset is inversely proportional to the length of the CAG repeat^6^. However, age of onset may be accelerated or delayed by genetic modifiers, including a SNP at the transcription factor binding site of Nf-kB in the *HTT* promoter^7^, and in genes related to DNA repair^8^.

The HTT protein is a large monomeric protein whose function is intricately regulated by post-translational modifications including phosphorylation, acetylation, ubiquitination, proteolysis, and fatty acylation^9^. While some PTMs of HTT have been shown to be protective against toxicity of mutant HTT, such as phosphorylation at S13/16 and S421^10^, others are crucial for HD pathogenesis or increase mutant HTT toxicity. In particular, caspase-mediated proteolysis of HTT at amino acid D586 has been shown to be necessary for the development of disease phenotypes in HD mouse models^11,12^. Consequently, modulating PTMs has become a focus of therapeutic strategies for HD. We sought to identify human SNPs that lead to missense mutations that may alter PTMs in HTT and, consequently, modify progression or pathogenic effects of the disease.

## Results

To identify SNPs that could alter HTT PTMs and potentially modify HTT function, all common missense mutations (≥0.1% minor allele frequency; MAF) within *HTT* were curated from Phase 3 of the 1000 Genomes Project (1KG) and from the Genome Aggregation Database (gnomAD). Nineteen common missense SNPs with ≥0.1% MAF were found in 1KG, and 19 common missense SNPs with ≥0.1% MAF were found in gnomAD (Table 1). The top 14 most common missense SNPs in 1KG and in gnomAD were shared in both data sets, highlighting convergent allele discovery by distinct methodologies.

**Table 1:** Functional SNPs in gnomAD and 1000 Genomes Phase 3.

| RSID | Chromosome<br>Position<br>(GRCh37.p13) | Reference | Alternate | Amino acid<br>position<br>(NP_002102.4) | Mutation | Allele Frequency<br>(gnomAD) | Allele<br>Frequency<br>(1000 G) | PROVEAN<sup>22</sup><br>PREDICTION | SIFT<sup>23</sup><br>PREDICTION | SNAP<sup>24</sup><br>PREDICTION |
| --- | --- | --- | --- | --- | --- | --- | --- | --- | --- | --- |
| rs362331 | 3215835 | T | C | 2311 | Y/H | 0.43680 | 0.44129 | Neutral | Tolerated | Neutral |
| rs362272 | 3234980 | G | A | 2788 | V/I | 0.28390 | 0.21026 | Neutral | Tolerated | Neutral |
| rs363125 | 3189547 | C | A | 1722 | T/N | 0.12750 | 0.18910 | Neutral | Tolerated | Neutral |
| rs149109767 | 3230410 | AGAG | A | 2645 | E/- | 0.06067 | 0.02656 | Deleterious | NA | ---- |
| rs35892913 | 3148570 | G | A | 1066 | V/I | 0.04200 | 0.02356 | Neutral | Tolerated | Neutral |
| rs363075 | 3137674 | G | A | 895 | G/R | 0.04069 | 0.02376 | Deleterious | Damaging | Effect |
| rs1143646 | 3148653 | T | G | 1093 | I/M | 0.03962 | 0.02236 | Neutral | Damaging | Neutral |
| rs3025837 | 3174845 | A | C | 1387 | N/H | 0.01489 | 0.04153 | Neutral | Tolerated | Neutral |
| rs34315806 | 3162034 | C | T | 1262 | T/M | 0.00837 | 0.02756 | Neutral | Damaging | Neutral |
| rs151106561 | 3132038 | A | G | 627 | Q/R | 0.00523 | 0.00280 | Neutral | Tolerated | Neutral |
| rs150003356 | 3189510 | T | C | 1710 | C/R | 0.00281 | 0.00100 | Deleterious | Tolerated | Effect |
| rs3025843 | 3156038 | A | G | 1175 | T/A | 0.00256 | 0.00679 | Neutral | Tolerated | Neutral |
| rs118005095 | 3129240 | G | A | 553 | G/E | 0.00242 | 0.00619 | Neutral | Tolerated | Neutral |
| rs34389685 | 3133113 | G | A | 698 | G/E | 0.00190 | 0.00200 | Deleterious | Damaging | Effect |
| rs186355914 | 3109095 | C | T | 233 | A/V | 0.00170 | 0.00020 | Neutral | Damaging | Neutral |
| rs201702168 | 3221998 | A | T | 2446 | E/D | 0.00154 | 0.00140 | Neutral | Tolerated | Neutral |
| rs375957195 | 3230428 | C | A | 2647 | D/E | 0.00153 | 0.00319 | Neutral | Tolerated | Neutral |
| rs1065746 | 3148624 | G | C | 1084 | D/H | 0.00135 | 0.00020 | Deleterious | Damaging | Effect |
| rs190593027 | 3144528 | C | G | 996 | T/R | 0.00110 | 0.00140 | Neutral | Damaging | Neutral |
| rs201646519 | 3210567 | T | C | 2076 | S/P | 0.00018 | 0.00100 | Neutral | Tolerated | Neutral |
| rs201760820 | 3134411 | G | A | 789 | V/M | 0.00047 | 0.00100 | Neutral | Tolerated | Neutral |

A manually curated list of known HTT PTMs was intersected with all common missense SNPs identified from 1KG and gnomAD (Figure 1). Among the 21 common missense SNPs identified from 1KG and gnomAD, two overlapped directly with known HTT PTMs; rs118005095 (G553E) and rs201646519 (S2076P) (Table 1, boxed; **Supplemental Table 1**). SNP rs201646519 intersects with the predicted phosphorylated residue S2076, resulting in a proline substitution thereby blocking phosphorylation at this site. Myristoylated G553^13,14^ overlapped with the missense SNP rs118005095, which leads to a substitution of glycine to glutamic acid (G553E) and would therefore completely abrogate post-translational myristoylation at this site (Figure 2a). Myristoylation involves the covalent addition of the 14 carbon fatty acid, myristate, to N-terminal glycines either co-translationally on the nascent polypeptide as it is being translated on the ribosome following the removal of the initiator methionine or post-translationally following proteolysis that exposes an N-terminal glycine^15^. The fatty acid moiety promotes membrane binding of proteins. No other common missense SNPs directly intersected with a known HTT PTM residue. Due to the low frequency of rs201646519, we focused on rs118005095.

**Figure 1.**
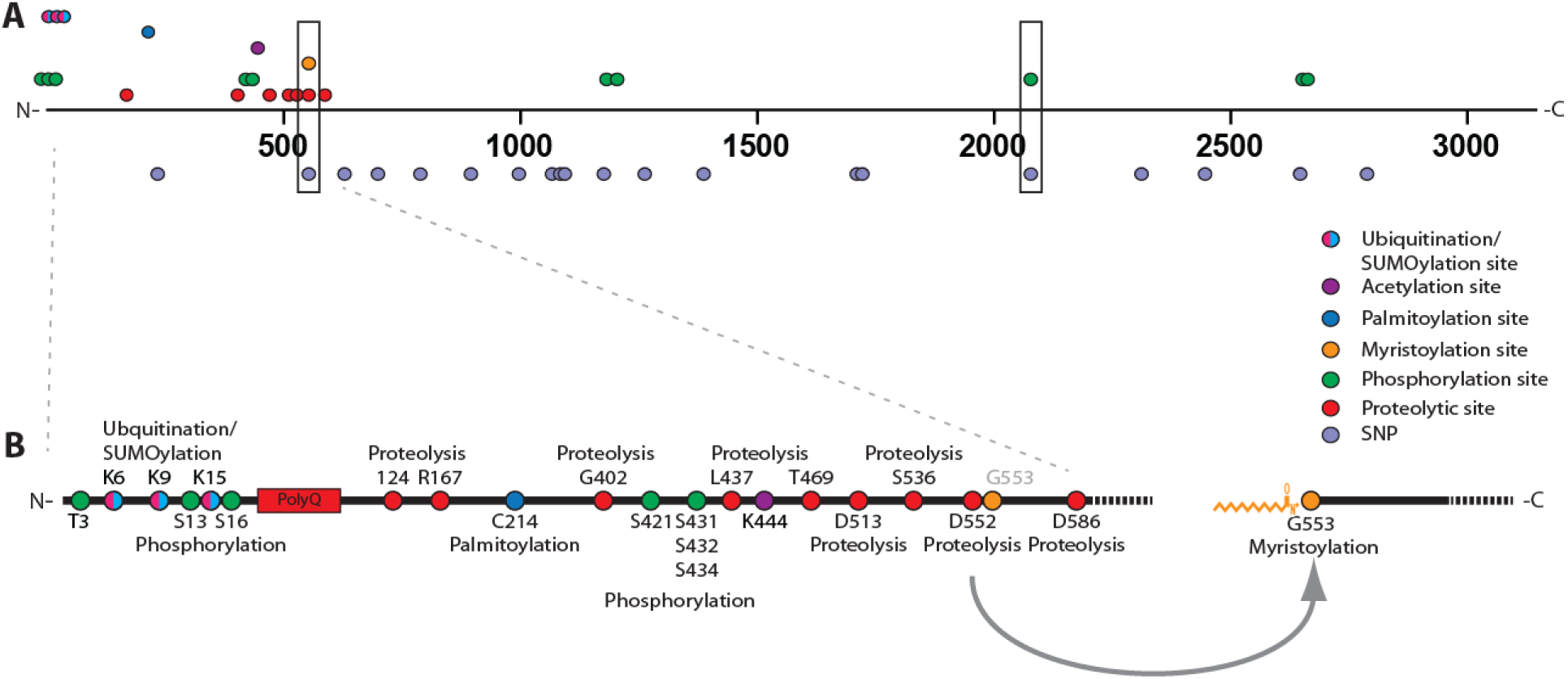
Schematic representing where HTT PTMs and *HTT* missense SNPs intersect. (A) Linear map of *HTT* SNPs leading to missense mutations mapped to known HTT PTMs. The two missense mutations that directly intersect with myristoylation at G553 and phosphorylation at S2076, G553E and S2076P, are boxed. (B) PTMs within the first 586 amino acids of HTT are highlighted. Proteolytic caspase sites are indicated on the bottom while non-caspase mediated proteolytic sites are displayed on top. G553 is myristoylated following caspase cleavage at D552.

**Figure 2.**
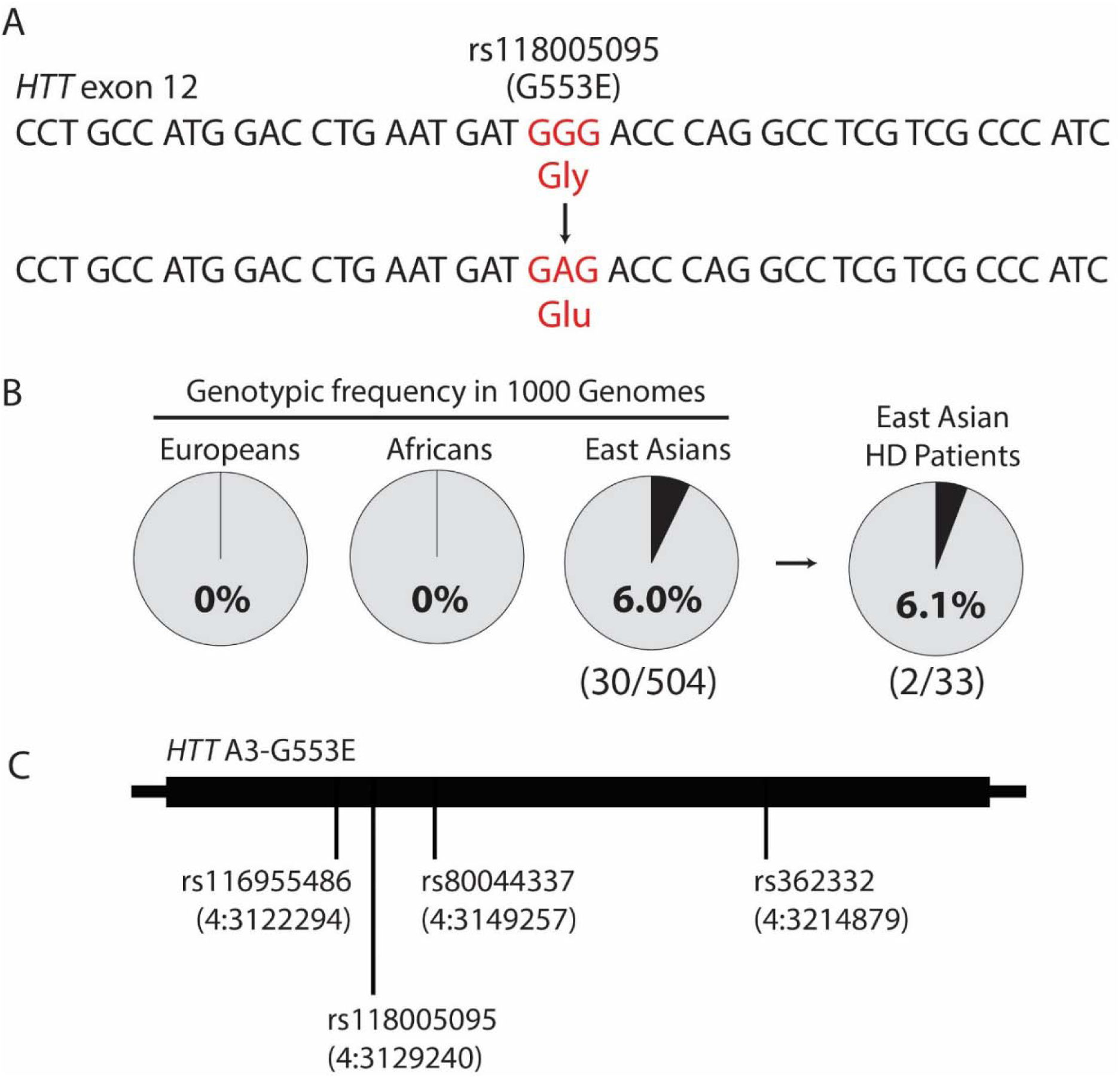
The rs118005095 missense variant is a naturally occurring human SNP that alters the HTT amino acid sequence. (A) rs118005095 results in mutation of the HTT 553 glycine residue to glutamic acid. (B) rs118005095 occurs exclusively in populations of East Asian ancestry. (C) The rs118005095 variant is one of four SNPs defining a gene-spanning *HTT* haplotype in the East Asian population.

The G553E SNP rs118005095 shows pronounced ethnic differences in frequency, being most common in individuals of East Asian ancestry. In 1KG, rs118005095 is observed on 31 out of 5008 chromosomes from all populations, of which 30 instances occur in subjects of defined East Asian origin (n=504 subjects) at a genotypic frequency of 6.0% (30/504) and allelic frequency of 3.0% (30/1008) (Figure 2b). The one remaining 1KG chromosome with rs118005095 outside East Asian individuals occurs in a Bengali subject from Bangladesh, close to East Asia. In the gnomAD data, rs118005095 occurs in chromosomes from East Asian subjects at an allelic frequency of 2.914% (552/18942). In contrast, rs118005095 is observed in <0.1% of chromosomes from European, African, South Asian, and Latino subjects, reflecting its absence in similar reference populations from 1KG. Therefore rs118005095 is expected to occur in approximately 6.0% of individuals from the East Asian general population.

We have previously shown that *HTT* is characterized by a haplotype block of low recombination and that SNPs within the gene represent specific haplotypes^16^. *HTT* haplotype analysis in 1KG reveals that rs118005095 occurs on a specific A3b haplotype variant in the East Asian population, tagged by three additional haplotype-defining SNPs: rs116955486, rs80044337, and rs362332 (Figure 2c). Of note, previously published haplotypes of the HD mutation in East Asian patients do not include this haplotype, suggesting that this SNP is not found on mutant *HTT* chromosomes in the East Asian population. We thus predicted that rs118005095 may occur in wild-type *HTT* of East Asian HD patients, potentially impacting the normal function of myristoylation in the context of the disease.

We have previously shown that the expanded polyglutamine mutation dramatically reduces myristoylation of HTT at G553^14^. HD patients with the G553E mutation (rs118005095) would therefore be unable to compensate for deficient myristoylation with a wild-type copy bearing the rs118005095 variant. Genotyping of archived DNA from 33 East Asian HD patients in the UBC HD Biobank confirmed the presence of the SNP in two patients (Figure 2b). As expected, phasing of rs118005095 to the mutation revealed that this missense variant occurred on wild-type HTT in these patients. However, no clinical information was available to assess the potential modifying effects of the SNP.

To investigate the predicted cellular effects of the rs118005095 (G553E) variant, we generated the mutation in a biologically relevant form of HTT bearing the first 1-588 amino acids appended to YFP (HTT_1-588^-^_YFP, Figure 3a), previously shown to be post-translationally myristoylated^14^ and disease-causing in HD mouse models^17^. Myristoylation at G553 was first validated in WT and mHTT_1-588^-^_YFP using an alkyne-myristate analog that can be covalently linked to biotin-azide via Click chemistry for detection (**Supplementary Figure 1**)^18^. Caspase-cleavage at D552 was promoted using staurosporine (STS) and cycloheximide (CHX) in order to increase post-translational myristoylation at G553, as previously described^14^. Post-translational myristoylation was detected in fully-cleavable (FC) HTT (HTT_1-588^-^_YFP with no point mutations added) even in the absence of STS/CHX, likely due to basal levels of caspase activity (**Supplementary Figure 1**). Myristoylation was increased in the presence of STS/CHX. A point mutation at the primary site of caspase-cleavage in HTT at position D586 to glutamate was made in order to more easily measure myristoylation at G553 following caspase cleavage at D552 (D586E, **Supplementary Figure 1**). This led to a dramatic increase of post-translational myristoylation at G553.

**Figure 3.**
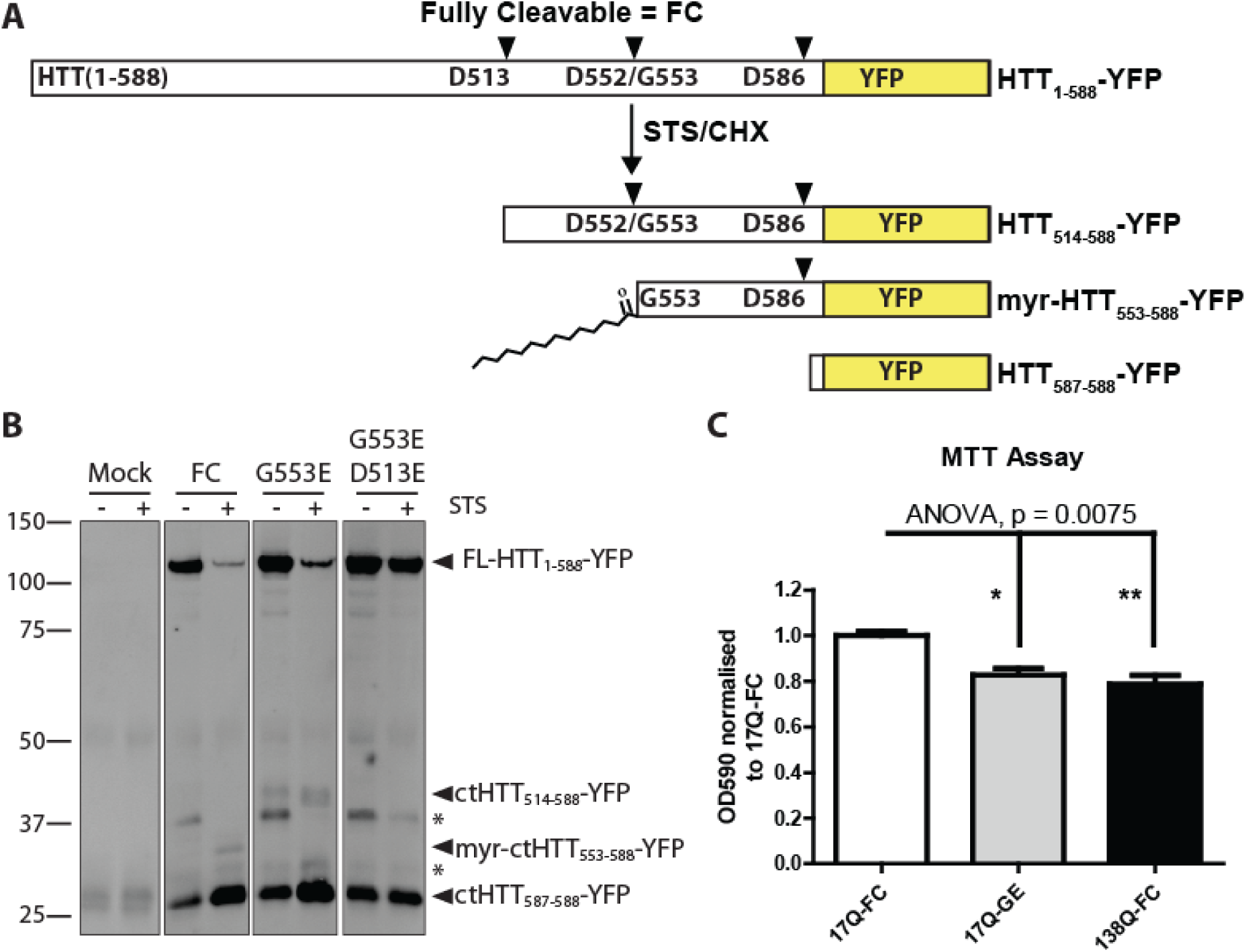
The G553E substitution blocks post-translational myristoylation and alters proteolysis of wtHTT leading to toxicity. (**A**) Schematic representation of proteolytic cleavage of WT fully cleavable (FC) HTT_1-588^-^_YFP. HTT is proteolysed by caspases at D586 and D552 and post-translationally myristoylated at G553. (**B**) HeLa cells were transfected with WT HTT_1-588^-^_YFP bearing the indicated point mutations. Proteolysis was induced with staurosporine (STS) and cycloheximide. In the presence of the G553E mutation, a higher molecular weight band was detected that was blocked when D513 was mutated to E, confirming the D513 caspase cleavage site. Similarly, the D586E mutation completely abrogated the generation of the ∼26 kDa band corresponding to caspase cleavage at D586. * denotes unknown or non-specific bands. (C) HeLa cells were transfected with the indicated constructs and cell viability was measured using the MTT assay (Kruskal-Wallis ANOVA, error bars represent SEM. *-p<0.05, ** - p<0.01).

As predicted, substitution of the essential glycine to alanine or serine (G553A/D586E and G553S/D586E) completely abrogated post-translational myristoylation of HTT at G553 following caspase-induced cleavage. Similarly, blocking caspase cleavage at D552 by substituting the aspartate to glutamate (D552E, **Supplementary Figure 1**) completely blocked post-translational myristoylation at G553. Inhibiting caspase-cleavage at D552 appeared to block the generation of both the myristoylated and non-myristoylated bands. Therefore, we determined that myristoylation status at G553 could also be determined by gel mobility. Non-myristoylated C-terminal (ct) HTT_553-588^-^_YFP migrates at a slightly higher molecular weight than myristoylated-ctHTT_553-588^-^_YFP (**Supplementary Figure 1**). This mobility shift was shown previously in a shorter myristoylated fragment ^14^and mimics other lipidated proteins such as LC3^19^.

Consequently, the G553E mutation was further characterized in the absence of the myristate analog in WT-HTT_1-588^-^_YFP (Figure 3). As expected, substitution of the essential N-terminal glycine at position 553 completely abrogated post-translational myristoylation of the wild-type form (Figure 3b). Surprisingly, the G553E mutation also inhibited caspase cleavage of HTT at D552, which was associated with a concomitant increase in caspase cleavage of HTT at D513 (Figure 3b). Cleavage at D586 did not appear to be affected. Of note, blocking myristoylation conventionally by substituting glycine to alanine did not prevent cleavage at D552 (**Supplemental Figure 1**). Proteolysis at D513 was recently shown to be toxic in cells ^11^. Due to the increased cleavage of HTT at the toxic site D513 in the presence of the G553E mutation, we hypothesized that the G553E variant may induce toxicity of wild-type HTT. Indeed, the G553E mutation significantly increased the toxicity of wild-type HTT to similar levels as mutant 138Q HTT alone (Figure 3c).

## Discussion

Previously, we have shown that proteolysis at D586 is essential to the pathogenesis of HD in a mouse model^12^. In addition, we identified post-translational myristoylation as a novel PTM in HTT that is decreased in the presence of the expanded polyQ suggesting it is a protective PTM^14^. Now, we have identified the first SNP leading to a missense mutation that alters multiple PTMs in HTT. This is also the first demonstration of a naturally occurring missense mutation that blocks post-translational myristoylation. The G553E mutation was shown to completely abrogate post-translational myristoylation of HTT and induce cellular toxicity of the protein in *cellulo*. Surprisingly, this mutation gives rise to a toxic fragment of wild-type HTT caused by proteolysis at D513.

We have shown the G553E mutation blocks caspase cleavage at D552 leading to increased cleavage of HTT at the toxic caspase site D513. It has recently been shown that proteolysis of either wild-type HTT or mutant HTT at more than one site, particularly caspase-cleavage at D513 and D586, is toxic^11^. This is mediated by a newly appreciated toxicity associated with the C-terminal HTT that is exacerbated by toxic shorter N-terminal fragments^11,20^. It is thought that the N- and C-terminal fragments interact in the presence of single site cleavage, which is considered less toxic. However, increased cleavage at multiple sites leads to decreased interaction following proteolysis at more than one site, particularly caspase cleavage at D513 and D586 and calpain cleavage at R167^11^. This proteolysis is toxic in both wild-type and mutant HTT.

Consequently, increased proteolysis of wild-type HTT at D513 caused by the G553E mutation, in conjunction with cleavage at D586 and loss of protective myristoylation, is predicted to be toxic. Loss of function of the mutant HTT allele would likely exacerbate this toxicity. Remarkably, blocking cleavage at D586 in an HD mouse model has been shown to completely ameliorate the disease phenotype in mice^12^. Therefore, we predict that the G553E mutation may be an important disease modifier and will lead to an earlier onset of disease in HD patients. It will be important to screen East Asian HD patients for the G553E mutation to validate the proteolytic consequences and phenotypic impact of the rs118005095 variant.

## Materials and Methods

### Genotyping and Haplotype Analysis

Haplotype analysis of rs118005095 in 1000 Genomes Phase 3 was performed using custom scripts in the R statistical computing environment applied to a variant call file spanning *HTT* variants (chr4:3034088-3288007) in all 5008 subjects^16^. SNPs in high pairwise linkage disequilibrium (r^2^>0.90) with rs118005095 were found in similarly high linkage disequilibrium on a common gene-spanning haplotype in subjects of East Asian ancestry among all populations in 1000 Genomes. Direct genotyping of rs118005095 was performed on banked DNA samples from East Asian HD subjects from the UBC HD Biobank. PCR amplicons containing the G553E variant (rs118005095) were generated using rs118005095 forward (GCGGACTCAGTGGATCTGGC) and rs118005095 reverse (GTCTGAAGGGGTAACAGCTGAATC) primers and Sanger sequenced using one or both primers.

### Cloning

All HTT-YFP point mutations were generated using gBlocks, as described previously^21^, except the D513E mutation was generated by Top Gene Technologies. All point mutations were confirmed by DNA sequencing (Operon Eurofins).

### Click chemistry and fragment analysis

Post-translational myristoylation of HTT_1-588^-^_YFP was detected as described previously^14^. Briefly, HeLa cells were transiently transfected using X-tremeGene (Roche) as per the manufacturer’s directions. After 20h post-transfection, the media was exchanged with DMEM containing charcoal filtered FBS and cells were incubated with alkyne-myristate (Cayman Chemicals) for 30 mins prior to the addition of 1 µM staurosporine (STS) and 5 µg/mL of cycloheximide (CHX) to promote caspase activity and prevent synthesis of new protein, respectively. After 4h, cells were lysed in modified RIPA buffer. HTT_1-588^-^_YFP was immunoprecipitated using goat anti-GFP (Eusera) and subjected to Click chemistry using azido-biotin (Invitrogen). Following SDS-PAGE on large 10% gels, post-translational myristoylation was detected by Western blot analysis using streptavidin Alexa680 (Invitrogen) and Licor. YFP was detected using rabbit anti-GFP (Eusera).

### MTT assay

Cell viability was measured using the MTT assay (Sigma) as per the manufacturer’s directions. Briefly, 20 h post-transfection. MTT was added to the cells for an additional 4 h. OD readings were normalized to WT fully cleavable HTT. The experiment was performed 6 times with 3 technical replicates per experiment for each condition.

## Acknowledgements

This work was supported by the CHDI Foundation and the Canadian Institutes of Health Research (CIHR 20R90174). DDOM was supported by postdoctoral fellowships from CIHR, Michael Smith Foundation for Health Research and the Bluma Tischler Fellowship from UBC. CK was supported by a CIHR Doctoral Research Award. The authors would like to thank Dr. Niels Skotte for contributing illustrations in Figure 1.

